# YY1 Binding to Regulatory Elements that Lack Enhancer Activity Promotes Locus Folding and Gene Activation

**DOI:** 10.1101/2023.08.23.554459

**Authors:** Zeqian Gao, Miao Wang, Alastair Smith, Joan Boyes

## Abstract

Enhancers activate their cognate promoters over huge distances but how enhancer/promoter interactions become established is not completely understood. There is strong evidence that cohesin-mediated loop extrusion is involved but this does not appear to be a universal mechanism. Here, we identify an element within the mouse immunoglobulin lambda (Igλ) light chain locus, HSCλ1, that has characteristics of active regulatory elements but lacks intrinsic enhancer or promoter activity. Remarkably, knock-out of the YY1 binding site from HSCλ1 reduces Igλ transcription significantly and disrupts enhancer/promoter interactions, even though these elements are >10 kb from HSCλ1. Genome-wide analyses of mouse embryonic stem cells identified 3503 similar YY1-bound, putative genome organizing elements that lie within CTCF/cohesin loop boundaries but that lack intrinsic enhancer activity. We suggest that such elements play a fundamental role in locus folding and in facilitating enhancer/promoter interactions.

**Highlights:** How long-range enhancer-promoter interactions are established is not fully understood An element in the lambda light chain locus, HSCλ1, lacks intrinsic enhancer activity Removal of YY1 binding from HSCλ1 disrupts neighbouring enhancer/promoter contacts Genome-wide analyses detect similar elements that lack enhancer or promoter activity We propose these elements aid locus folding and nearby enhancer-promoter interactions

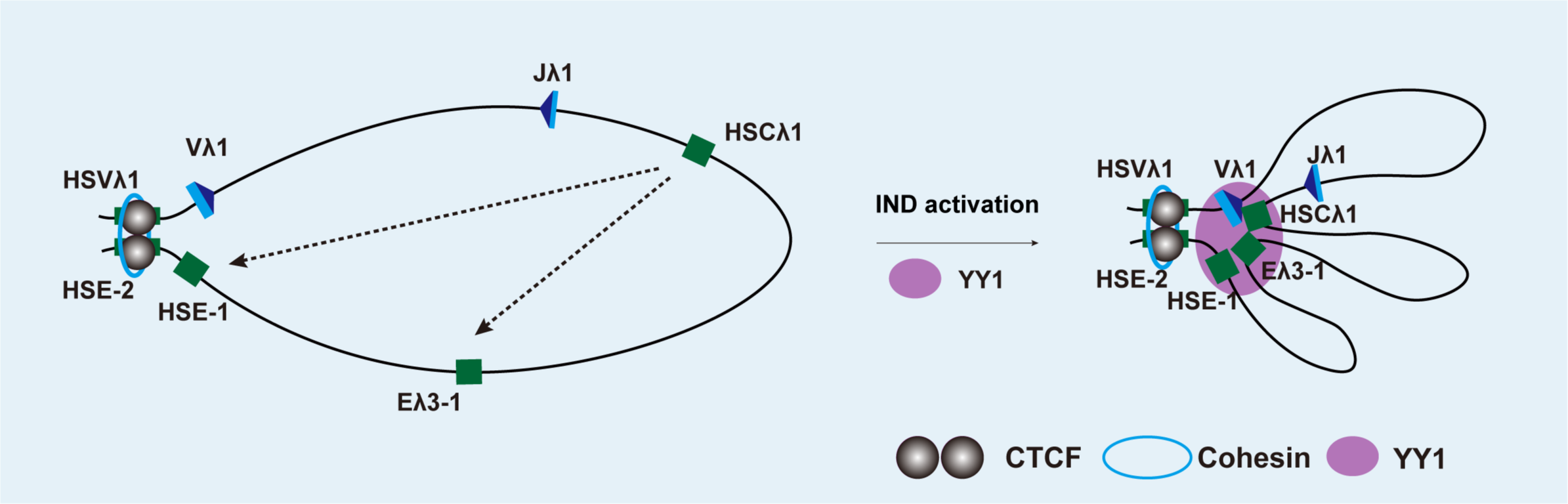

## Introduction

Enhancers play a pivotal role in driving spatiotemporal gene regulation by activating target gene expression over distances of ∼2 kb to >2 Mb. This long-range activation is underpinned by the arrangement of chromatin into a series of topologically associating domains (TADs) or insulated neighbourhood domains (INDs), where the chromatin loop boundaries are typically anchored by CTCF/cohesin [1]. Enhancer/promoter interactions primarily take place within INDs where enhancers physically contact their cognate promoter via long range chromatin looping, as demonstrated by chromatin conformation capture, and derivative experiments [2, 3]. Enhancer/promoter loops are then stabilised via transcription activators including Yin Yang 1 (YY1) and Mediator such that Mediator physically links transcription factors at enhancers with promoter-bound transcription machinery [4]. By contrast, YY1 binds to enhancers or promoters via either its DNA or RNA binding domains and forms homodimers that can bridge enhancer/promoter loops [5]. Genome wide analyses found that in most cases, YY1 binding connects active enhancers and promoters in enhancer/enhancer, enhancer/promoter and promoter/promoter interactions, although a small fraction of YY1 is associated with insulators in both mouse embryonic stem cells (mESCs) and mammalian cell lines [5]. Consistent with its functional role in stabilising enhancer/promoter loops, ablation of YY1 binding sites within enhancers eliminated enhancer/promoter interactions whereas artificial tethering of YY1 to enhancers, restored these interactions [5].

Initial formation of enhancer/promoter loops is thought to be achieved by cohesin-mediated loop extrusion that brings enhancers and promoters into close proximity. However, complete depletion of cohesin has only a small effect on gene regulation [6-8]. This may be because the depletion experiments were carried after most enhancer/promoter interactions had been established [9] and consistent with this, the effects of cohesin depletion were most apparent on enhancer activation of inducible genes [10]. It was also found in experiments using the same enhancer/promoter pair, that interactions over large distances (>100 kb) are more dependent on cohesin than those that span <50 kb [11]. Similar studies in cortical neurons showed that cohesin depletion affected enhancer/promoter interactions over >1.5 Mb but had a much lower impact on those of <40 kb [12]. Given this, and the fact that cohesin depletion preferentially effects genes close to CTCF/cohesin loop boundaries [13], it seems likely that additional mechanisms operate to establish enhancer/promoter contacts.

The murine immunoglobulin lambda light chain locus (Igλ) is an ideal model to study how such interactions might be brought about due to its relatively small size (230 kb) and the presence of only six functional V and J gene promoters and 3-4 regulatory elements. The locus appears to have been duplicated during evolution and available Hi-C data suggest that CTCF/cohesin form insulated neighbourhood domains in the 5’ and 3’ halves of the Igλ locus by the early pro-B cell stage [14]. Activation of the Igλ locus occurs considerably later at the pro-B/pre-B transition when transcription of the V and J gene segments is upregulated by ∼8-fold [15]. A fundamental question therefore is how enhancer/promoter interactions are established at the pro-B/pre-B transition within the pre-formed CTCF/cohesin loops. We identified an element, HSCλ1 that shows many of the characteristics of active enhancers and lies between known Igλ enhancers and promoters but lacks binding sites for critical activators. Given that it has a binding site for YY1, it seems possible that this element facilitates locus folding that then promotes enhancer/promoter interactions.

Here, we investigated the role of this element and show that it indeed lacks intrinsic enhancer activity but that YY1/HSCλ1 binding is vital for long range locus folding, adjacent enhancer/promoter interactions and crucially, for full levels of locus transcription. Complementary genome-wide analyses in mESCs identified YY1 binding to 3503 similar putative genome organising elements that also lack intrinsic promoter and enhancer activity, as determined by self-transcribing active regulatory region sequencing (STARR-seq) [16]. Analyses of individual loci in mESCs imply that the putative genome organising elements function similarly to HSCλ1. These studies therefore identify a new class of genome organising elements that appear integral to locus folding and enhancer/promoter interactions.

## Results

Using available ChIP-seq data from early B cells, we previously identified two new enhancer-like elements, HSE-1 and HSCλ1 in the 3’ half of the Igλ locus, in addition to the well characterised Eλ3-1 enhancer that has been shown to activate Jλ1 transcription [17-19] (Figure 1(A)). Of the new enhancer-like elements, HSE-1 is very similar to Eλ3-1 in that it binds identical transcription factors [14], including E2A and PU.1/IRF4 that are key activators of immunoglobulin light chain loci [18, 20]. Formation of a CTCF/cohesin-bound loop between HSVλ1 and HSE-2 (Figure 1 (A)), that lie at the boundaries of the Igλ 3’ domain, will bring HSE-1 into closer proximity of Vλ1. To test if this might allow HSE-1 to enhance Vλ1 transcription, we first mapped the Vλ1 promoter using 5’-RACE; this identified transcriptional start sites (TSS) at ∼39 bp upstream of the Vλ1 coding sequence (N=6). A 650 bp sequence that maps to the ATAC-seq peak spanning the Vλ1 TSS [14] was then cloned into a luciferase reporter construct with or without HSE-1 or HSCλ-1. Consistent with its predicted enhancer activity, the presence of HSE-1 increases Vλ1 transcription by >12-fold compared to the promoter-only construct following transfection into the pre-B cell line, 103/BCL-2 [21] (Figure 1(B)). By contrast, HSCλ-1 lacks a PU.1 binding site [14] and consequently, IRF4, that shows only weak independent DNA binding [22], displays only low occupancy of HSCλ1 [14]. As can be seen in Figure 1(B), HSCλ1 causes only negligible increases in Vλ1 transcription following transfection of luciferase reporter constructs into 103/BCL-2 cells [21].

**Figure 1:**
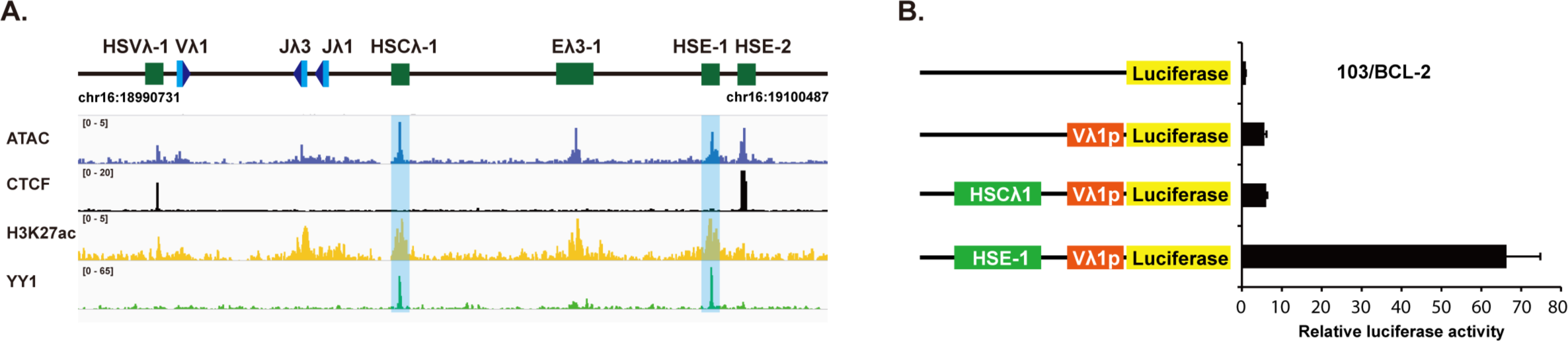
The enhancer, HSE-1, activates Vλ1 transcription, in contrast to HSCλ1. A. Upper: Schematic of the murine Igλ locus. Putative regulatory elements are depicted by green rectangles; V and J gene segments by light blue rectangles and recombination signal sequences by dark blue triangles. Lower: ATAC-seq and ChIP seq data mapped to the Igλ locus; all data are from pre-B cells. B. Luciferase activity driven by the Vλ1 promoter ±HSE-1 or HSCλ1 enhancer in 103/BCL-2 cells that had been temperature shifted to 39.5°C [21]. The Vλ1 promoter increases luciferase activity by ∼5-fold compared to the empty vector; HSE-1 gives a further >12-fold increase whereas HSCλ1 has only a negligible effect. Error bars show standard error of the mean (SEM) from three biological replicates.

HSCλ1, however, has characteristics that are typical of active enhancers including open chromatin, enrichment of H3K27 acetylation, as well as occupancy by some activators such as p300, E2A, Mediator [14] and notably, YY1 [2]. HSCλ1 is located between the Eλ3-1 enhancer and the Jλ1/Jλ3 promoters (Figure 1(A)); given that YY1 is central to long range interactions [5], we hypothesised that HSCλ1 might be involved in locus folding and/or in facilitating enhancer/promoter interactions. To test this idea, we specifically deleted the YY1 binding site in HSCλ1 using CRISPR/Cas9 (Figure 2(A)) in PIPER-15 cells, a pro-B cell line in which Igλ transcription can be induced [14] and which is therefore a good model system to investigate how enhancer/promoter interactions are established at the pro-B/pre-B transition. Even though HSCλ1 has no enhancer activity *per se*, removal of the YY1 binding site in HSCλ1 significantly reduced Vλ1 transcription both prior to induction (basal transcription) and following induction (Figure 2(B)). To determine if this is due to altered locus folding, 3C analyses were performed using Eλ3-1 as the viewpoint. These showed a significant reduction in interactions between the Eλ3-1 enhancer and HSCλ1, as might be expected if YY1 bridges the two enhancer-like elements. Remarkably, however, interactions between Eλ3-1 and Vλ1 and Jλ1 promoters, that are >20 kb and >10 kb from HSCλ1, respectively, are also significantly reduced. By contrast, interactions between the Eλ3-1 and HSE-1 enhancers that are 90 kb apart but where YY1 sites remain intact, are unaffected (Figure 2(C)). These data therefore imply that HSCλ1 is important in locus folding and notably, also for interactions between enhancers and promoters that are distant from HSCλ1 itself. Consistent with this idea, interactions between HSCλ1 and the enhancer elements, HSE-1 and Eλ3-1, appears to result in the Vλ1 and Jλ1 promoters being brought into closer proximity of the enhancer hub (Graphical abstract).

**Figure 2:**
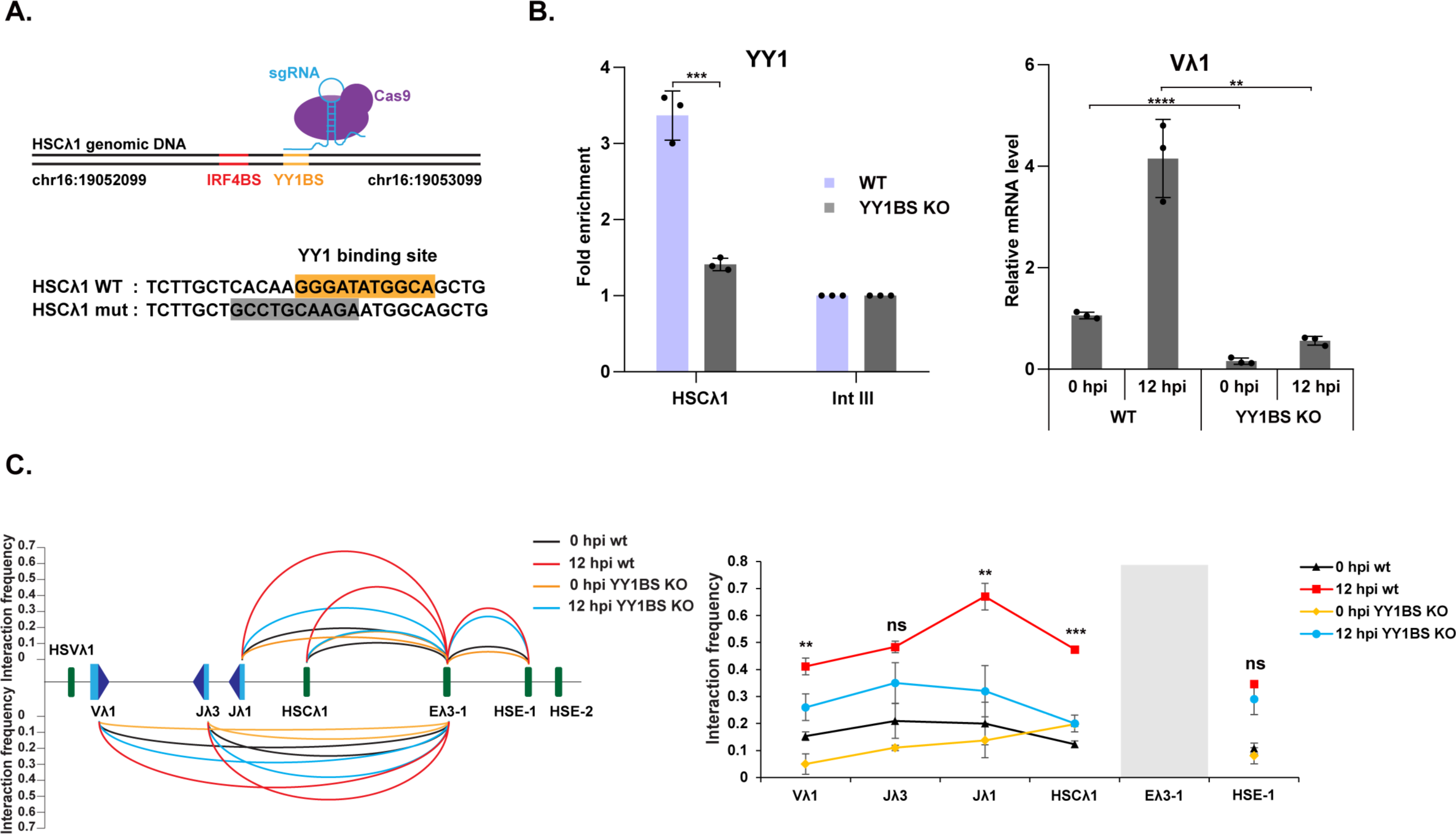
Effects of knock-out of YY1 site in HSCλ1. A. Upper: Schematic of targeting of YY1 binding site within HSCλ1 by CRISPR/Cas9. Lower: Sequence alignments of a clone bearing a deletion of the YY1 binding site at HSCλ1 with the wild type (WT) sequence. B. Left: YY1 binding was analysed by ChIP-qPCR in wild type (WT) PIPER-15 cells and cells where the binding site at HSCλ1 has been knocked-out (YY1BS KO). The fold enrichment over input DNA at HSCλ1 and Intgene III is shown. All values are normalized to binding at Intgene III as a negative control. Right: RT-qPCR analysis of Vλ1 transcription in cells where the YY1 binding site is knocked-out. Data are normalized to *Hprt* expression. C. Left: Analysis of the relative interaction frequency of Dpn II fragments from the Eλ3-1 viewpoint in induced wild type and YY1 binding site knockout PIPER-15 cells. The height of curves between Eλ3-1 and other genomic fragments represents the average value of interaction frequency obtained from three experimental repeats (Supplementary Table 1). Data were normalized using an interaction within the *Ercc3* locus. The plots to the right show the significance of the difference in interactions at 12 hours post induction between wild type and YY1 binding site knockout cells. Error bars show standard error of the mean (SEM) from three biological replicates.

The requirement to establish enhancer/promoter interactions via mechanisms that are distinct from cohesin-mediated loop extrusion is more broadly relevant and we were keen to determine if YY1-bound, putative genome organising (YGO) elements, like HSCλ1, are a more general phenomenon. To this end, we capitalised on available genome-wide data from mESCs that used self-transcribing active regulatory region sequencing (STARR-seq) to identify all elements with enhancer activity. Here, randomly sheared DNA is placed downstream of a minimal promoter driving GFP and upstream of a poly A site. Active enhancers trigger upregulation of their own transcription and can therefore be identified by reverse transcription and sequencing the RNA products [16]. We combined these data [23] with available ChIP-seq and capture Hi-C data from mESCs to identify YY1-bound elements within INDs that lack intrinsic enhancer activity. In these analyses, firstly, INDs were identified by integrative analysis of capture Hi-C and CTCF ChIP-seq data [24]. Whole genome YY1 binding sites were then intersected with the INDs to obtain intra-IND YY1 binding elements. YY1 binds to various regulatory elements, including promoters, enhancers and IND boundaries [5]. The intra-IND YY1 elements which can be mapped to TSS ±2 kb, enhancers ±2 kb, as identified by STARR-seq [23], and left or right boundaries ±2 kb were discarded to obtain Intra-IND YY1 binding elements without intrinsic enhancer activity. From this, we identified 3503 elements that have high YY1 occupancy and lie in open chromatin but lack enhancer activity. To further investigate the characteristics of these elements, we plotted ChIP-seq signals of frequently observed, active histone modifications (H3K27ac and H3K4me3) and of transcription factors (CTCF, cohesin, YY1, Nanog, RNAPII and Mediator) to the putative genome organising elements as well as to other DNA regulatory elements within active INDs. As shown in Figure 3, active gene promoters show high levels of ATAC-seq signals, H3K4me3 signals, Mediator and RNAPII binding, whereas YY1-bound enhancers (YBEnh) show high levels of STARR signals, ATAC signals, H3K27ac signals, Mediator and Nanog binding. Notably, the YY1-bound putative genome organising elements (YGOs) show moderate levels of histone modifications and transcription factor binding but high YY1 occupancy, suggesting YGOs are a novel type of DNA regulatory element that is distinct from normal enhancers.

**Figure 3:**
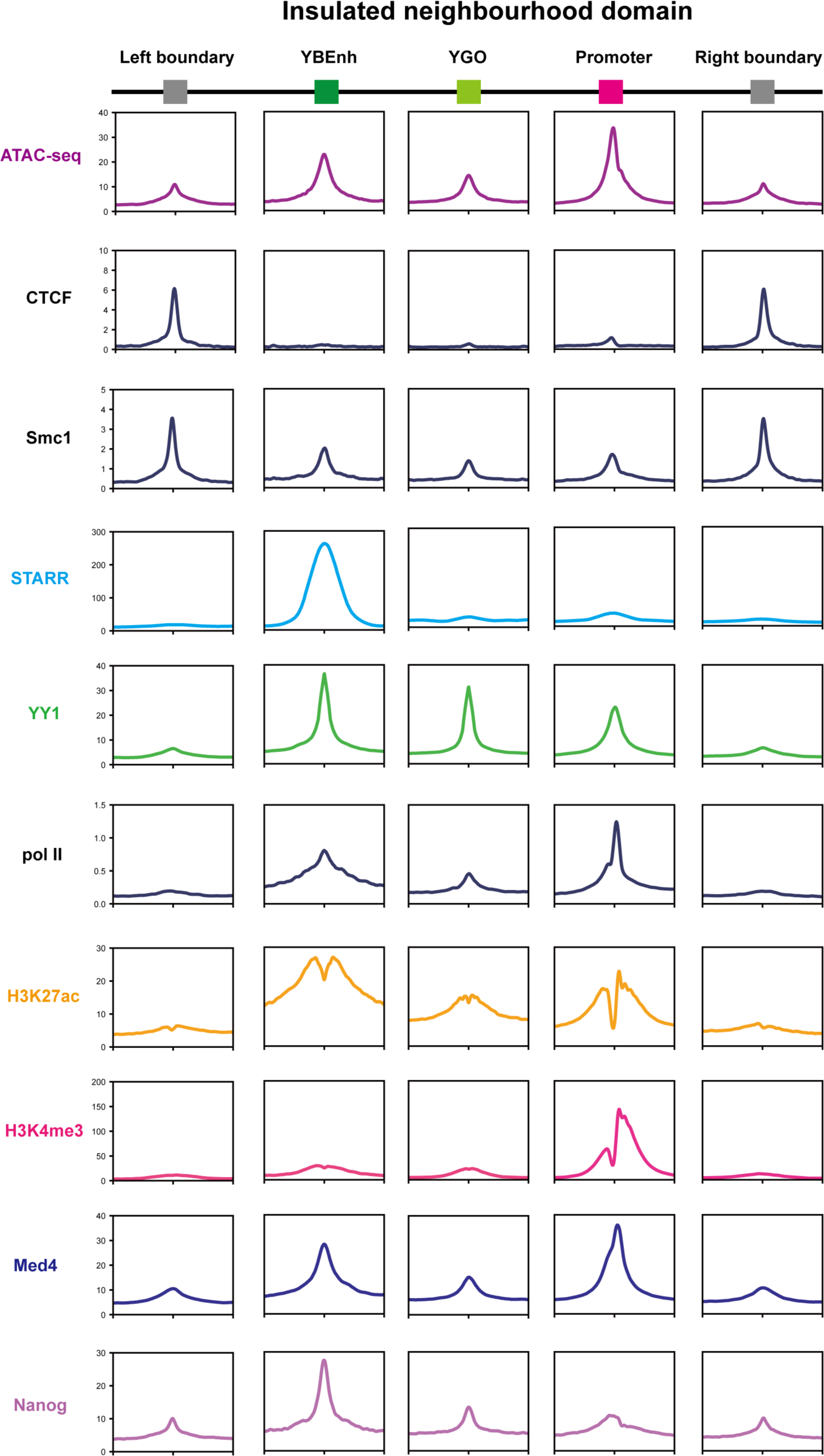
Genome-wide analyses in mESCs of YY1-bound elements that lack intrinsic enhancer activity. Schematic of active INDs that have YGOs and YY1-bound enhancers (YBEnh) in E14 mESCs. The genome coordinate for each IND were obtained from Atlasi et al [26]. The genomic coordinates of IntraIND gene promoters and YBEnh were obtained by intersecting INDs with H3K4me3 peaks and STARR peaks, respectively. The genome coordinates of YGOs were obtained by removing YY1 peaks mapped to YBEnh and gene promoters from IntraIND YY1 sites. Normalized reads for STARR-seq and ATAC-seq data as well as ChIP-seq data for H3K27ac, H3K4me3, CTCF, Smc1, YY1, Nanog, Med4, RNAPII were plotted to the genome coordinates of the IND boundaries, YGOs, YBEnh and gene promoters, respectively.

To further investigate the function of YGOs in mESCs, we next mapped ChIP-seq and STARR-seq data to two distinct INDs on chromosome 3. This identified YGOs within each IND with high YY1 occupancy but moderate levels of histone modifications and transcription factor binding. As can be seen in Figure 4, the positions of the YGOs varies between INDs but in each case the YGO has the potential to impact locus folding and enhancer/promoter interactions. These studies therefore imply that YY1-bound putative genome organising elements are a general phenomenon.

**Figure 4:**
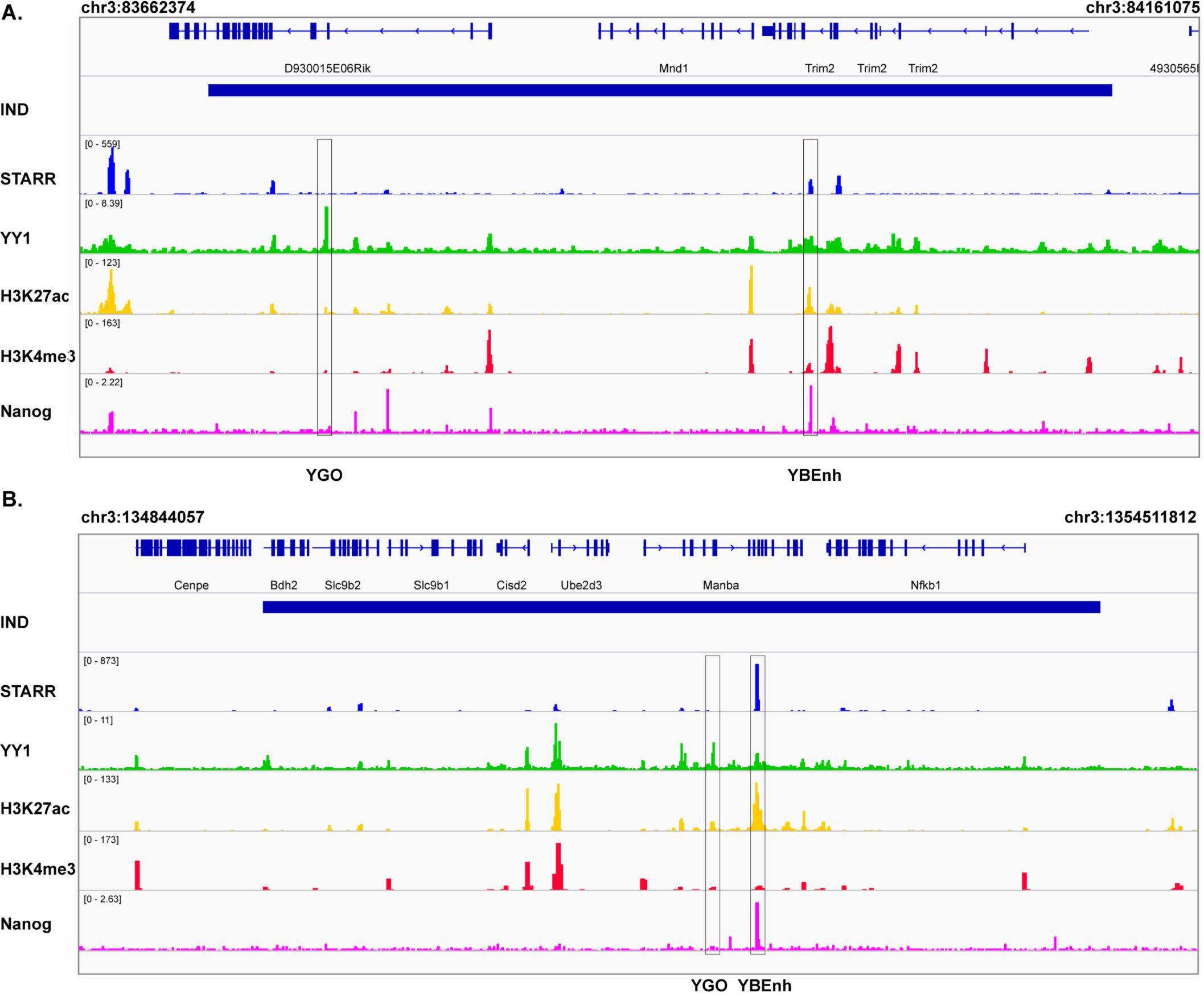
Putative Genome Organising elements are present in active INDs in mESCs. A. Characterisation of elements in the IND at chromosome 3:83662374-83161075 in E14 mESCs. The blue bar represents an active IND (spanning ∼500 kb) which was identified using capture Hi-C and CTCF ChIP-seq data from E14 mESC. Active promoters, at the genes mapped at the top of the Figure, show increased H3K4me3 whereas the putative YY1-bound genome organising element, labelled YGO, is located at 5’ half of the IND and shows high YY1 occupancy but no enhancer activity, determined by STARR-seq (STARR). A YY1-bound enhancer (YBEnh) is located in the 3’ half of the IND and shows high levels of H3K27Ac, YY1 and Nanog binding. B. Characterisation of elements in the IND at chromosome 3:134844057-135451812 in E14 mESCs. As for (A); in this case, the YGO and YBEnh are located in the middle of the IND.

## Discussion

Enhancers physically interact with their cognate promoters, in some cases over huge distances, within the tightly packed eukaryotic nucleus but the mechanism by which enhancer/promoter contacts are initially established remains incompletely understood. Cohesin-mediated loop extrusion is an attractive mechanism that can explain how many, but not all, long-range interactions become established [9]. Here, we provide evidence for a putative new type of regulatory element, namely YY1-bound genome organising elements (YGO), that appear to mediate localised chromatin folding within INDs to facilitate enhancer/promoter interactions. Remarkably, removal of the YY1 binding site from HSCλ1 causes a profound reduction in Vλ1 transcription even though HSCλ1 lacks intrinsic enhancer activity. Furthermore, there is a dramatic alteration in 3C interactions, including a significant reduction in Eλ3-1/Vλ1 and Eλ3-1/Jλ1 interactions even though these elements are >10 kb from HSCλ1. Given that YY1 is known to homodimerize and to stabilise long-range genome interactions, these data suggest that YY1 binding to HSCλ1 alters locus folding, leading to increased interactions of enhancers and promoters within the insulated neighbourhood domain. Consistent with its important functional role, HSCλ1 is present in the 3’ half of the duplicated Igλ locus but an equivalent element is not found in the 5’ half of the locus. Remarkably, Vλ1 recombination, in the 3’ half of the Igλ locus, accounts for ∼70% of Igλ recombination events compared to only 15% for the Vλ2 and Vλx gene segments in the 5’ domain [25] even though the recombination signal sequences adjacent to Vλ1 and Vλ2 are identical.

Our genome-wide analyses in mouse ESCs support the idea that elements with properties like HSCλ1 are a more general phenomenon. Currently, mESCs are the only cell type where both STARR-seq and ChIP-seq data are available and thus where intrinsic enhancer activity has been measured genome-wide. Nonetheless, we identified 3503 YGOs. The position of these elements varies between loci but it is notable that these lay within INDs and analyses of individual loci suggest that their locations are consistent with a role in promoting interactions between enhancers and promoters. Whilst our *in silico* analyses can’t exclude the alternative possibility that some of the elements function as cohesin loading sites to facilitate loop extrusion, in this case, it would be difficult to explain the highly significant binding of YY1. Notably, the elements we identify in mammalian cells share functional characteristics with the tethering elements that were recently identified in *Drosophila melanogaster* [26]. Tethering elements also have characteristics in common with active enhancers such as H3K4 mono-methylation and binding of transcription activators, although binding of YY1 was not noted. These elements promote enhancer-promoter interactions independent of TAD boundaries and given that 620 tethering elements were identified compared to 2034 insulators (putative TAD boundaries), the authors proposed that tethering elements form a complementary mechanism of genome organisation to TADs [26]. We identify 3503 YY1-bound elements (compared to ∼15,000 INDS in mammalian nuclei [24]) that are found in open chromatin and show characteristics of moderately active enhancers (H3K27 acetylation and binding of activators) but lack intrinsic enhancer activity; it seems possible that these elements organise the mammalian genome in a similar way to which tethering elements organise the Drosophila genome.

YY1 has properties that make it well suited to mediating long range interactions, including its ability to bind both DNA and RNA, to form homodimers and to interact with intrinsically disordered domains in other transcription factors [5]. Consequently, it has the potential to be tethered to enhancers via enhancer RNAs [27], interact with different transcription factors at promoters and to homodimerize with other YY1 proteins that bind to DNA sequences within other regulatory elements. Consistent with a critical role for YY1 in stabilising enhancer/promoter interactions, published studies show that removal of YY1 binding sites ablated enhancer/promoter interactions whereas inducible degradation of YY1 over 24 hours led to significant changes in expression of >8000 genes [5]. By contrast, recent studies have shown that depletion of YY1 has only a marginal effect on enhancer/promoter interactions genome-wide [28]. These depletion experiments, however, were performed over only over three hours and it is possible that this short time-scale was insufficient to trigger changes in genome organisation.

Overall, our studies suggest that YY1 binding to non-enhancer, non-promoter, non-insulator sites maintains functional chromatin domains to facilitate locus folding and associated enhancer/promoter interactions.

## Materials and methods

### Vectors

LentiCRISPR v2 was obtained from AddGene (#52961) and was a kind gift of Feng Zhang whereas pCMVR8.74 and pMD2.G (#22036 & #12259) were kind gifts of Didier Trono. pGL3-Vλ1p was constructed by cloning the Vλ1 promoter (chr16: 19084509-19085159) in front of luciferase reporter gene in pGL3-Basic (Promega). To construct pGL3-Vλ1p-HSE-1 and pGL3-Vλ1p-HSCλ1, HSE-1 (chr16: 19007329-19008187) or HSCλ1 (chr16: 19026931-19027772) were cloned ∼3 kb upstream of the Vλ1 promoter in pGL3-Vλ1p. Signal guide (sg)RNAs targeting HSCλ1 were designed using the online design software (http://crispr.mit.edu) and cloned into the lentiCRISPR v2.

### Cell lines

HEK293T were maintained in Dulbecco’s Modified Eagle Medium (DMEM) supplemented with 10% foetal calf serum, 4 mM L-glutamine, 50 U/ml penicillin and 50 μg/ml streptomycin. Cells were grown in a humified incubator at 37°C with 5% CO_2_.

103/BCL-2 [21], a kind gift from Prof. Naomi Rosenberg) and PIPER-15 cells [14] were maintained at a density of 0.5-2 x 10^6^ cells /ml, in complete Roswell Park Memorial Institute (RPMI)-1640 medium supplemented with 10% foetal calf serum, 4 mM L-glutamine, 50 U/ml penicillin and 50 μg/ml streptomycin and 50 μM β-mercaptoethanol. Cells were grown at 33°C with 5% CO_2_. PIPER-15 cells were induced to activate Igλ transcription by addition of Tamoxifen to a final concentration of 2 μM [14].

### Transfection of 103/BCL-2 cells and Luciferase reporter assays

Electroporation was carried out using the Nucleofector^TM^ Kit (LONZA # VPA-1010) according to manufacturer’s instructions and as further detailed in [14]. Luciferase assays were carried out using the Dual-Luciferase Kit (Promega) according to manufacturer’s instructions and as detailed in [14].

### Production of lentiviral particles

Lentiviral particles were produced in HEK293T cells by transfection with the lentiviral backbone constructs, packaging construct (pCMVR8.74) and envelope construct (pMD2.G). 3 x 10^6^ HEK293T cells were plated per 10 cm dish in complete DMEM 24 hours before transfection. Three hours prior to transfection, the medium was changed to DMEM supplemented with 5% foetal calf serum, 4 mM L-glutamine. Separately, 4.9 μg of lentiCRISPRv2, 2.6 μg of pCMVR8.74 and 2.5 μg of pMD2.G were mixed with 500 μl of OptiMEM medium by gentle vortexing; in parallel, 30 μl of PEI stock solution (1 mg/ml) was diluted with 500 μl of OptiMEM medium. The solutions were then mixed by gentle vortexing for 15 seconds, followed by incubation at room temperature for 15 minutes and dropwise addition to cells. Cells were incubated at 37°C for 48 and 72 hours prior to harvest. The lentivirus-containing supernatant was filtered through a 0.45 μm syringe filter, flash frozen on dry ice and stored at -80°C until use.

### Knockout of the YY1 site in HSCλ1

CRISPR sgRNA oligonucleotides that target the YY1 site in HSCλ1 (Supplementary Table 1) were designed as described above. The YY1HSCλ1_sgRNA oligonucleotides were annealed and cloned into lenti-CRISPR v2 and used to produce lentiviruses. To transduce PIPER-15 cells, 5 x 10^5^ cells were spin-fected with 500 μl of sgRNA lentivirus and selected with 0.25 μg/ml puromycin after 48 hours. After one week of selection, monoclonal cell lines were generated using semi-solid agar; clones were screened for knockouts by PCR using the primers HSCλ1delR and HSCλ1delF (Supplementary Table 1). Monoclonal cell lines with apparent deletions in these regions were amplified using the above primers; the products were cloned and knockout of the respective region confirmed by Sanger sequencing.

### Chromatin Immunoprecipitation (ChIP)

ChIP experiments were performed according to Nowak et al [29] by first cross-linking with 2 mM Disuccinimidyl Glutarate (DSG, Sigma 80424) and then with 1% formaldehyde. The anti-YY1 antibody (22156-1-AP; Proteintech) was used at the dilution recommended by the manufacturer. The recovered DNA was analysed using quantitative PCR and the primers shown in Supplementary Table 1.

### Chromatin conformation capture (3C)

3C was carried out according to Dekker et al [30] with modifications. These, and the preparation of control BAC template, are described [14]. A nested PCR assay to detect 3C interactions was performed as described [14] using Eλ3-1 as the viewpoint and the primers given in [14]. Nested PCR reactions were also performed on the BAC control template to correct for differences in primer efficiency. The first round of PCR was performed using Taq DNA polymerase. For the second round, TaqMan qPCR was conducted in duplicate in 10 μl final volume with 5 μl of 1:10 diluted first round PCR product, 400 pM each primer, 100 pM 5’ nuclease probe and 5 μl qPCRBIO probe mix (PCRBIO PB20.21-05). All 3C samples were normalised by analysis of interactions in the *Ercc3* locus which is expected to be consistent across all cell types [31].

### Total RNA extraction and reverse transcription

Total RNA was extracted from approximately 2 x 10^6^ cells using TRIzol (Invitrogen #3289) according to the manufacturer’s instructions, followed by treatment with 2 U DNase I, as described [14]. 1 µg of RNA was reverse transcribed with M-MuLV reverse transcriptase (Invitrogen) as described [14].

### 5’ Rapid Amplification of Complementary DNA Ends (5’-RACE)

This was performed according to “Rapid amplification of 5’ cDNA ends” [32] with modifications. RNA (1 µg) was reverse transcribed as above and the oligo dT primer was removed by the addition of three volumes of buffer QG (Qiagen) and one volume isopropanol before application to a Qiagen quickspin column (Qiagen) and elution according to manufacturer’s instructions. To generate A tailed cDNA, the extracted cDNA was then added to 1 x Terminal Transferase buffer (NEB), 250 µM CoCl_2_ (NEB), 100 µM dATP, 10 U Terminal Transferase (NEB) and ddH_2_O to a volume of 50 µl. The tailing reaction was performed at 37°C for 30 minutes after which the enzyme was inactivated by heating at 75°C for 15 minutes and tailing reaction components were diluted by increasing the volume to 100 µl with ddH_2_O.

To generate 5’-RACE products, the A-tailed cDNA was subjected to PCR in a reaction that comprised 5-20 µl diluted A-tailed cDNA (∼50 ng assuming 1:1 RNA to cDNA conversion), 1 x Q5 Reaction buffer (NEB), 200 nM dT adaptor primer and *Vλ1* specific primer (Supplementary Table 1), 200 µM dNTPs and 2.5 U Q5 Hot-Start polymerase (NEB). A touchdown protocol was used to increase the specificity of the PCR; the thermal profile consisted of 98°C for 3 minutes, followed by 15 cycles of 98°C for 10 s, 71°C for 20 s and 72°C for 1.5 minutes, 10 cycles of 98°C for 10 s, 68°C for 20 s and 72°C for 1.5 minutes, 15 cycles of 98°C for 10 s, 65°C for 20 s and 72°C for 1.5 minutes and a final extension at 72°C for 3 minutes.

The highest intensity bands were excised and gel extracted. These products were eluted in 30 µl ddH_2_O and 1 µl was used in a PCR reaction designed to add a Hind III restriction site to the 3’ end of the product, to enable cohesive end cloning (a Xho I recognition site was present in the dT Adaptor primer). The PCR reaction consisted of 1x ThermoPol Buffer (NEB), 200 µM dNTPs, 200 nM dT adaptor primer and 200 nM Vλ1-GSP4-2-Hind III and 2 U Taq polymerase (NEB), in a final volume of 50 µl. The thermal profile was: 94°C for 3 minutes followed by four cycles of 94°C for 30 s, 58°C for 20 s, and 68°C for 2 minutes and 16 cycles of 94°C for 30 s, 60°C for 20 s, 68°C for 2 minutes with a final extension at 68°C for seven minutes. 5’-RACE products were purified by phenol-chloroform extraction and ethanol precipitation, cloned into pBluescript SK- and Sanger sequenced.

### Real-time PCR using SYBR Green

Quantitative PCR was performed using a Corbett Rotor-Gene 6000 machine and analysed using the Corbett Rotor-Gene 6000 Series Software (v.1.7, build 87). A typical qPCR reaction contained 5 μl 2×SensiFAST SYBR No-Rox mix (Bioline #BIO-98080), 2∼10 ng DNA template, or cDNA at a final dilution of 1:100, 400 nM of each primer in a total volume of 10 μl. Primer sequences are given in Supplementary Table 1. All reactions were performed in duplicate. In each case, a standard curve of the amplicon was analysed concurrently to evaluate the amplification efficiency and to calculate the relative amount of amplicon in unknown samples. A melt curve, to determine amplicon purity, was produced by analysis of fluorescence as the temperature was increased from 72°C to 95°C.

### Analysis of next generation sequencing data

Accession numbers of all datasets used for this study are given in Supplementary Table 1. All datasets are from E14 mESCs, cultured in serum plus leukaemia inhibitory factor (LIF). ChIP-seq and ATAC-seq data were analysed as described previously [14]. STARR-seq peaks and YY1 binding peaks from E14 mESCs were called with MACS2 by Peng et al [23] and Atlasi et al [24], respectively. The details of defining INDs using capture HiC and CTCF ChIP-seq data from E14 mESCs are given by Atlasi et al [24]. IntraIND YGOs were obtained via removing YY1 peaks that mapped to IND boundaries ±2 kb, TSS ±2 kb and STARR enhancers ±2 kb using bedtools v2.18. The density of reads across each type of DNA regulatory element was plotted using deepTools 2.0 with default parameters. More specifically, the signal distribution of each type of DNA regulatory elements was computed using computeMatrix in the reference-point mode; plotHeatmap was used to generate the density map.

### Identification of Intra-IND YY1 binding elements without intrinsic enhancer activity

Initially, INDs were identified integrative analysis of capture Hi-C and CTCF ChIP-seq data [24]. Whole genome YY1 binding sites from ChIP-seq data were then intersected with INDs to obtain intra-IND YY1 binding elements. The intra-IND YY1 elements which mapped to TSS ±2 kb, STARR enhancers ±2 kb, left or right boundaries ±2 kb were discarded to obtain intra-IND YY1 binding elements without intrinsic enhancer activity.

## Statistical Analyses

Statistical analyses were performed using GraphPad Prism v9. Analyses of fold changes between biological replicates, using biologically distinct samples from the same types of cells, were performed using a paired Student’s *t* test where *p < 0.05, **p < 0.01, ***p < 0.001, ****p < 0.0001.

## Acknowledgements

This work was supported by China Scholarship Council studentships (to ZG and MW) and a studentship from the National Centre for the Replacement, Refinement and Reduction of Animals in Research (NC3Rs; NC/K001639/1 to ALS). We are very grateful to Prof. Naomi Rosenberg (Tufts University) for 103/BCL-2 cells.

## Declaration of interests

None

## Author contributions

ZG – Conceptulisation, Investigation, Data curation, Writing – review and editing.

MW - Investigation, Data curation, Writing – review and editing

ALS - Investigation, Data curation, Writing – review and editing

JB – Funding acquisition, Supervision, Project administration, Writing – original draft, Writing – review and editing.

## Notes

### Competing Interest Statement

The authors have declared no competing interest.

